# A kinome-wide synthetic lethal CRISPR/Cas9 screen reveals that mTOR inhibition prevents adaptive resistance to CDK4/CDK6 blockade in HNSCC

**DOI:** 10.1101/2023.08.07.552216

**Authors:** Yusuke Goto, Keiichi Koshizuka, Toshinori Ando, Hiroki Izumi, Xingyu Wu, Kyle Ford, Xiaodong Feng, Zhiyong Wang, Nadia Arang, Michael M. Allevato, Ayush Kishore, Prashant Mali, J. Silvio Gutkind

**Affiliations:** Moores Cancer Center, University of California San Diego, La Jolla, CA 92093, USA; Graduate School of Biomedical & Health Sciences, Hiroshima University, Japan; Department of Bioengineering, University of California San Diego, San Diego, CA 92093, USA

**Keywords:** CRISPR screen, synthetic lethality, HNSCC, signal transduction, cell cycle regulation mTOR inhibitor, palbociclib

## Abstract

The comprehensive genomic analysis of the head and neck cancer (HNSCC) oncogenome revealed frequent loss of p16^INK4A^ (*CDKN2A*) in most HPV negative HNSCC lesions, often concomitant with amplification of the cyclin D1 (*CCND1*) gene locus. However, cyclin-dependent kinase 4 and 6 (CDK4/6) inhibitors as single agents have shown modest effect in the clinic, even when combined with cetuximab. The aberrant activation of PI3K/mTOR pathway is highly prevalent in HNSCC, and recent clinical trials targeting mTOR showed promising results in terms of objective responses and progression free survival. However, the clinical efficacy of mTOR inhibitors (mTORi) for advanced HNSCC patients may be limited due to intrinsic or acquired resistance. By a kinome-wide CRISPR/Cas9 screen, we identified cell cycle inhibition as a synthetic lethal target for mTORi. Combination of mTORi and palbociclib, a CDK4/6 specific inhibitor, showed strong synergism in HNSCC-derived cells in vitro and in vivo. Remarkably, we found that adaptive increase in cyclin E1 (CCNE1) expression upon palbociclib treatment underlies the rapid acquired resistance to this CDK4/6 inhibitor in HNSCC. Mechanistically, mTORi inhibits the formation of eIF4G-*CCNE1* mRNA complexes, with the consequent reduction in mRNA translation and CCNE1 protein expression. Our findings suggest that concomitant mTOR blockade reverts the adaptive resistance to palbociclib, thereby providing a novel multimodal therapeutic option for HNSCC patients by co-targeting mTOR and CDK4/6. Our findings may have broad implications to halt the emergence of palbociclib resistance.

## Introduction

Head and neck squamous carcinoma (HNSCC) is among the ten most frequent cancers in the United States, with 54,540 new cases and 11,580 deaths estimated in the US alone in 2023 (1). Recent breakthrough treatment options by the use of immunotherapies targeting immune checkpoints brought survival benefit for HNSCC patients; however, the overall response rate to these immunotherapies in HNSCC is only ∼20% (2). Thus, novel therapeutic options for this disease are urgently needed.

The comprehensive analysis of the HNSCC oncogenome revealed frequent loss of p16^INK4A^ (*CDKN2A*) in HPV negative HNSCC, which account for 60% of the cases (3). Furthermore, amplification of the cyclin D1 (*CCND1*) gene is a frequent event in HNSCC, which has been reported to include 31 % of the HPV negative HNSCC (3). However, cyclin-dependent kinase 4 and 6 (CDK4/6) inhibitors as single agents have shown modest effects regardless of *CDKN2A-* altered status in recurrent and metastatic HNSCC (4). Furthermore, a double-blind, randomized phase II trial (PALATINUS) that evaluated the efficacy of palbociclib plus cetuximab in patients with unselected HPV-unrelated recurrent or metastatic HNSCC did not significantly prolong the overall survival (OS) of patients with HNSCC (5). In this context, novel combinatory therapeutics for palbociclib is in needed. Our group has been focusing on the study of mTOR signaling in HNSCC. Indeed, we have shown that the PI3K-mTOR pathway is the most frequently activated signaling mechanism in HNSCC, as judged by strong pS6 expression in more than 90% of HNSCC specimens (6). Based on these results, we have recently performed a clinical trial using rapamycin, which is a 1^st^ generation mTOR inhibitor (mTORi), in newly diagnosed HNSCC patients. Here, we found that rapamycin was effective for most of the HNSCC patients, with an overall response rate of 25 % including one case of complete response despite 21 days treatment duration (7). Similarly, we have recently shown that mTOR inhibition with everolimus diminish significantly the progression free survival of locally advanced HPV negative HNSCC lesions in the adjuvant setting (8). However, earlier clinical trials involving advanced, recurrent metastatic HNSCC patients showed limited response and resulted in treatment failure (9). The molecular mechanisms underlying mTORi resistance should be uncovered to find precise molecular targets which can be combined with mTORi to achieve durable responses.

In this study, we aimed to discover synthetic lethal targets and resistance mechanisms for mTORi by taking advantage of CRISPR/Cas9 screening. Using 2^nd^ generation mTORi, INK128, we identified cell cycle regulation pathway as one of most significant synthetic lethal targets and resistant pathways to mTORi in HNSCC. We confirmed strong synergism between INK128 and palbociclib, which is a widely used approved CDK4/6 inhibitor, *in vitro* and *in vivo*. In turn, we found that CCND1 and Cyclin E1 (CCNE1) accumulate upon palbociclib treatment, and that CCNE1 overexpression is sufficient to induce palbociclib resistance in HNSCC cells. Co-administration of INK128 and palbociclib could prevent the protein accumulation of CCNE1 by reducing mRNA translation, and consequently, co-administration of these targeted agents can revert the resistance to palbociclib in CCNE1 overexpressing cells. Overall, our findings suggest that co-targeting mTOR and cell cycle signaling represents a potential therapeutic option for HNSCC. These findings may be also relevant for other cancer types characterized by the progressive acquisition of resistance to CDK4/6 inhibitors.

## Results

### CRISPR/Cas9 screening identifies cell cycle regulation as a synthetic lethal mechanism for mTORi in HNSCC

To explore synthetic lethal targets and resistance mechanisms for mTORi in HNSCC, we took advantage of CRISPR screening. First, we generated Cas9-expressing Cal27 HNSCC cells (Cal27-Cas9) (Supp. Fig. 1A) and confirmed cutting efficiency using two different gRNAs (gT1/ gT2) targeting AAVS locus. Next generation sequencing (NGS) for these cells showed 83.0% and 98.3% of non-homologous end joining (NHEJ) frequency for gT1 and gT2, respectively (Supp. Fig. 1B), indicating the cutting efficiency for Cal27-Cas9 was suitable to conduct the planned screening. Since our purpose was to discover druggable targets, and the kinome is the target of a large proportion of oncology-related drugs, we used a human kinome-wide CRISPR library, targeting 763 genes consisting of 4 gRNAs for each gene (10). After infecting Cal27-Cas9 cells with the kinome-wide CRISPR library, we treated Cal27-Cas9-kinome cells with vehicle or mTORi until total population doubling reached 20 (Fig.1A and 1B). In this study, we applied INK128 (also known as MLN-0128 and TAK-228), which is an mTOR ATP-competitive small molecule inhibitor, and reported to have excellent physiochemical properties (11,12). After extracting DNA from these cells, we performed PCR to amplify the barcodes, and NGS to identify depleted gRNAs in mTORi-treated cells compared with vehicle-treated cells (Fig. 1A). remarkably, the most depleted gene was mTOR, which is consistent with the fact that we used a low dose mTORi to find synthetic lethal targets (Supplementary Table 1). We next applied pathway analysis for hit gRNAs not only focusing on single gRNAs but investigating hit gRNAs as an integrated set of genes to find effective pathways to target. KEGG pathway analysis for significantly depleted 109 gRNAs showed enrichment of ErbB signaling pathway, MAPK signaling pathway, and Cell cycle pathways (Fig. 1C). Co-targeting ErbB signaling with mTORi for HNSCC has been shown in several studies, including our combination study of cetuximab and rapamycin or everolimus for HNSCC (13). Also, our group has reported RNAi screen for HNSCC cells with first generation mTORi, rapamycin, and showed MAPK signaling as synthetic lethal pathway for rapamycin (14). These studies are consistent with the results from this CRISPR screening. From clinical data, the high mRNA expression or protein expression of Cyclin D1, which is encoded by the *CCND1* gene, plays a central role in cell cycle, and is a worse prognostic factor for overall survival in HNSCC patients (Fig. 1D and 1E), supporting an important role of cell cycle signaling for HNSCC. These results prompted us to explore the possibility of co-targeting cell cycle mechanisms and mTOR signaling in HNSCC.

**Figure 1.**
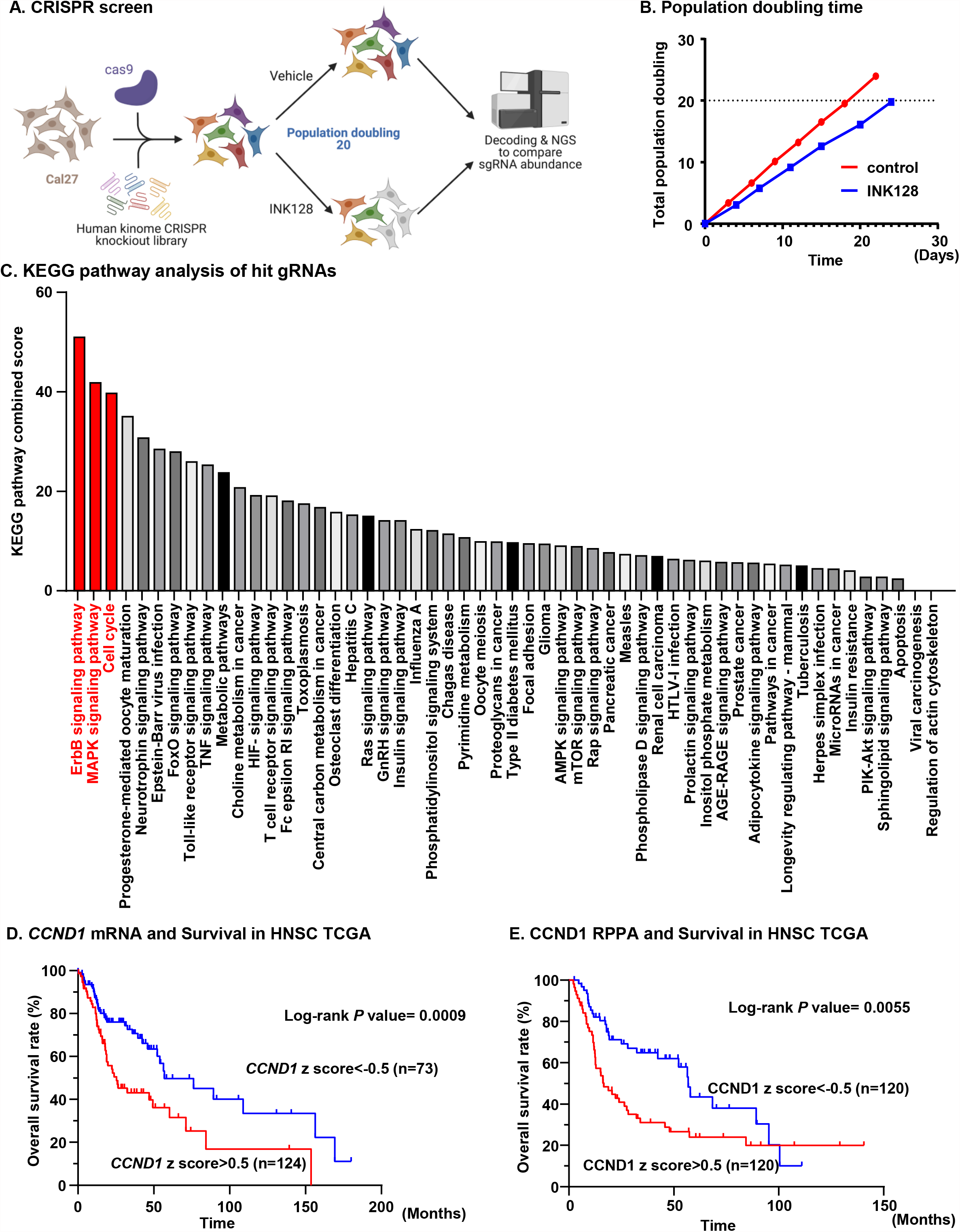
CRISPR screening identified cell cycle pathway as synthetic lethal pathway for mTORi in HNSCC. A, Scheme for CRISPR screening. Cal27-Cas9 cells were infected with human kinome CRIPSR knockout library, targeting 763 genes consisting of 4 gRNAs for each gene, and subjected to vehicle treatment or INK128 treatment. At population doubling (PD) 20, genomic DNA was extracted from cells, and PCR and NGS were performed. B, Total PD difference between two groups. At day 18, control cells proliferated to total PD of 20, and INK128 treated cells proliferated to total PD 20 at day24. C, KEGG pathway analysis for depleted gRNAs. KEGG pathway analysis was applied for significantly depleted 109 gRNAs in INK128 treated cells compared with control cells. ErbB signaling pathway, MAPK signaling pathway, and Cell cycle pathway were enriched. D, *CCND1* mRNA expression and overall survival in TCGA-HNSC patients. Patients with higher mRNA expression of *CCND1* (z score > 0.5; N = 124) demonstrated worse prognosis than those with lower mRNA expression (z score < -0.5; N = 73). *P* = 0.0009 (Log-rank test). E, CCND1 protein expression and overall survival in TCGA-HNSC patients. Patients with higher protein expression of CCND1 (z score > 0.5; N = 120) demonstrated worse prognosis than those with lower protein expression (z score < -0.5; N = 120). *P* = 0.0055 (Log-rank test).

### Combination of INK128 and palbociclib shows strong synergism in HNSCC cells *in vitro*

We next investigated the efficacy of blockade of both mTOR and cell cycle signaling. We took advantage of multiple cell lines, including Cal27 and HN12 cells, reflecting the human HPV negative HNSCC oncogenome (15). As an FDA-approved cell cycle targeting agent, we chose palbociclib, which can inhibit CDK4/CDK6 specifically. INK128 potently blocked the cell viability with growth inhibition of 50% (GI^50^) of 42 nM for Cal27 HNSCC cells (Fig. 2A). Similarly, GI^50^ for palbociclib was 1.27 μM for Cal27 cells (Fig. 2B). Next, we investigated the synergism between these drugs by the Chou-Talalay method (16). Fraction affected combination index (CI) plot showed CI of below 1 for most percentage of fraction when cells were treated with 1:10 or 1:20 concentrations of INK128 and palbociclib, respectively (Fig. 2C). These data suggest strong synergistic effect of this combination. Also, we performed a factorial dose matrix combinatorial drug treatment with INK128 and palbociclib, supporting synergism for this combination (Fig. 2D). Furthermore, we analyzed synergism using Bliss model (17), which suggested strong synergism with relatively higher INK128 concentration than GI^50^ (Fig. 2E). Next, to confirm the synergism in another cell line, we used HN12 HNSCC cells. The GI50 for INK128 in HN12 was 28 nM, and 0.85 μM for palbociclib (Fig. 2F and 2G). The combination index was below 1 when used 1:10 or 1:20 concentration of INK128 and palbociclib respectively, similar as Cal27 cells (Fig. 2H). To analyze the combination effect in conditions which are more reflective of cell growth in 3D, *in vivo* conditions, we tested orosphere assays which allows for the propagation of cancer cells that retained stemness and self-renewal (18). INK128 and palbociclib significantly reduced the number of sphere formation in Cal27 and HN12 cells, and co-administration of these two drugs could significantly block the sphere formation (Fig. 2I, J). These data suggest the possibility of using the combination of INK128 and palbociclib for treatment of HNSCC.

**Figure 2.**
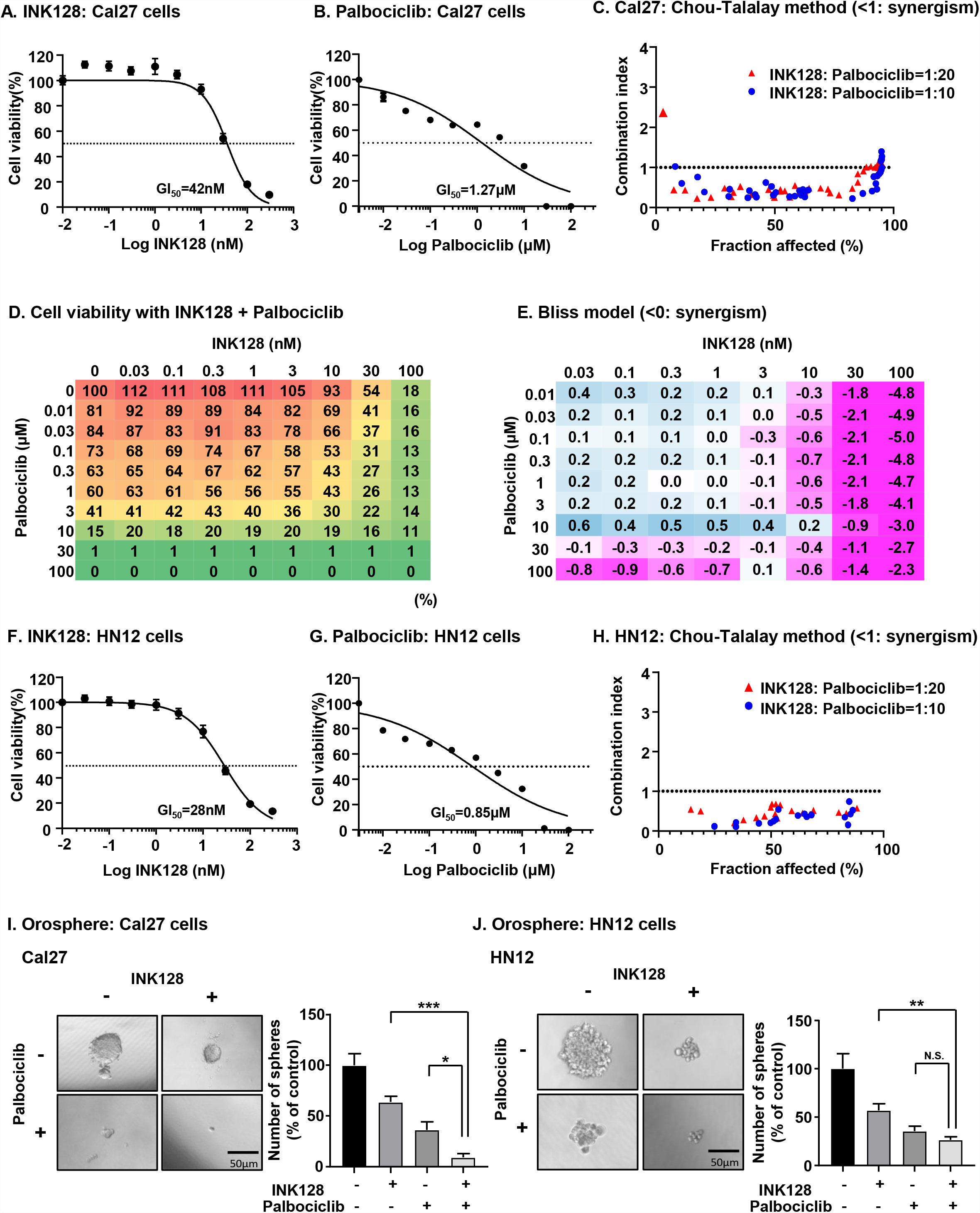
Combination of INK128 and palbociclib showed strong synergism in HNSCC cells in vitro. A, The effect of INK128 on Cal27 HNSCC cells. INK128 potently blocked the cell viability with growth inhibition of 50% (GI^50^) of 42 nM for Cal27 HNSCC cells. Bars represent average ± SEM (N = 3). B, The effect of palbociclib on Cal27 HNSCC cells. GI^50^ for palbociclib was 0.89 μM for Cal27 cells. Bars represent average ± SEM (N = 3). C, Analysis for synergism between INK128 and palbociclib by Chou-Talalay method for Cal27 cells. CI was below 1 for most percentage of fraction when cells were treated with 1:10 or 1:20 concentration of INK128 and palbociclib, respectively. D, Factorial dose matrix for INK128 and palbociclib (N = 3). E, Analysis for synergism between INK128 and palbociclib by Bliss model. Strong synergism was observed with relatively higher INK128 concentration than GI50. F, The effect of INK128 on HN12 HNSCC cells. GI^50^ for INK128 was 28 nM for HN12 HNSCC cells. Bars represent average ± SEM (N = 3). G, The effect of palbociclib on HN12 HNSCC cells. GI^50^ for palbociclib was 0.32 μM for HN12 cells. Bars represent average ± SEM (N = 3). H, Analysis for synergism between INK128 and palbociclib by Chou-Talalay method for HN12 cells. CI was below 1 for most percentage of fraction when cells were treated with 1:10 or 1:20 concentration of INK128 and palbociclib, respectively. I, Orosphere assay for Cal27 cells. The combination treatment significantly reduced the number of spheres compared with single agent of INK128 or palbociclib. Bars represent average + SEM (N = 5). \**P* < 0.05, \*\*\**P* < 0.001 (one-way ANOVA). J, Orosphere assay for HN12 cells. The combination treatment significantly reduced the number of spheres compared with single agent of INK128. Bars represent average + SEM (N = 5). \*\**P* < 0.01 (one-way ANOVA).

### Upregulation of CCNE1 by palbociclib confers resistance to palbociclib, which can be reverted by INK128

To investigate the mechanism for the synergism between INK128 and palbociclib in HNSCC, we explored changes in signaling components and cell cycle mechanisms. As expected, INK128 could effectively inhibit PI3K/ mTOR activation as judged by pAKT and pS6 expression levels and Cal27 and HN12 HNSCC cells (Fig. 3A). As for cell cycle, palbociclib treatment prevented phosphorylation of retinoblastoma protein (Rb), and caused upregulation of CCND1 and CCNE1. We hypothesized CCND1 and CCNE1 overexpression could represent a mechanism for palbociclib resistance in HNSCC, considering recent clinical data showing high *CCNE1* as worse clinical outcome for palbociclib treated patients in breast cancer (19). In this regard, we engineered HNSCC cell lines which stably overexpress CCND1 and CCNE1 individually, and together (Fig. 3B). This approach revealed increased resistance to palbociclib in Cal27-CCNE1 cells compared with Cal27-wt cells. However, no significant resistance to palbociclib was observed in Cal27-CCND1 cells. CCND1/E1-overxpressing Cal27 cells were also resistant, however, the resistance was similar as CCNE1-Cal27 cells, which suggests that no additional resistance was conferred by CCND1 overexpression (Fig. 3C). Similar results were confirmed using HN12-wt, HN12-CCND1, HN12-CCNE1, and HN12-CCND1/E1 and palbociclib (Fig. 3D). These data indicate that CCNE1 overexpression may represent one of the mechanisms of resistance to palbociclib in HNSCC. Remarkably, although we observed upregulation of CCNE1 after treatment with palbociclib in HNSCC cells, the addition of INK128 together with palbociclib could revert this overexpression and downregulated CCNE1 (Fig. 3A). Furthermore, the resistance of Cal27-CCNE1 and HN12-CCNE1 to palbociclib in terms of cell viability was completely abolished by addition of INK128 (Fig. 3E and 3F). Blockade of mTOR with INK128 has been shown to lead to the de-phosphorylation of 4E-BP1, and in turn, to the reduced binding between eIF4E and eIF4G, resulting in reduced mRNA translation of proliferative proteins (11). Indeed, INK128 treatment reduced eIF4G binding of the mRNA for *CCND1* and *CCNE1* (Fig. 3G and Supp. Fig. 1C). We hypothesized INK128 could revert the overexpression of CCNE1 caused by palbociclib treatment by reducing binding of eIF4G and *CCNE1*. As shown in Fig. 3G, combination treatment with INK128 and palbociclib potently reduced *CCNE1* mRNA binding to eIF4G compared with control or palbociclib treatment. These data suggest that CCNE1 overexpression is one of the resistance mechanisms to palbociclib, and that mTOR acts upstream of CCNE1, controlling its mRNA translation. Together, these data provide a rationale for the combination therapy of INK128 and palbociclib for HNSCC.

**Figure 3.**
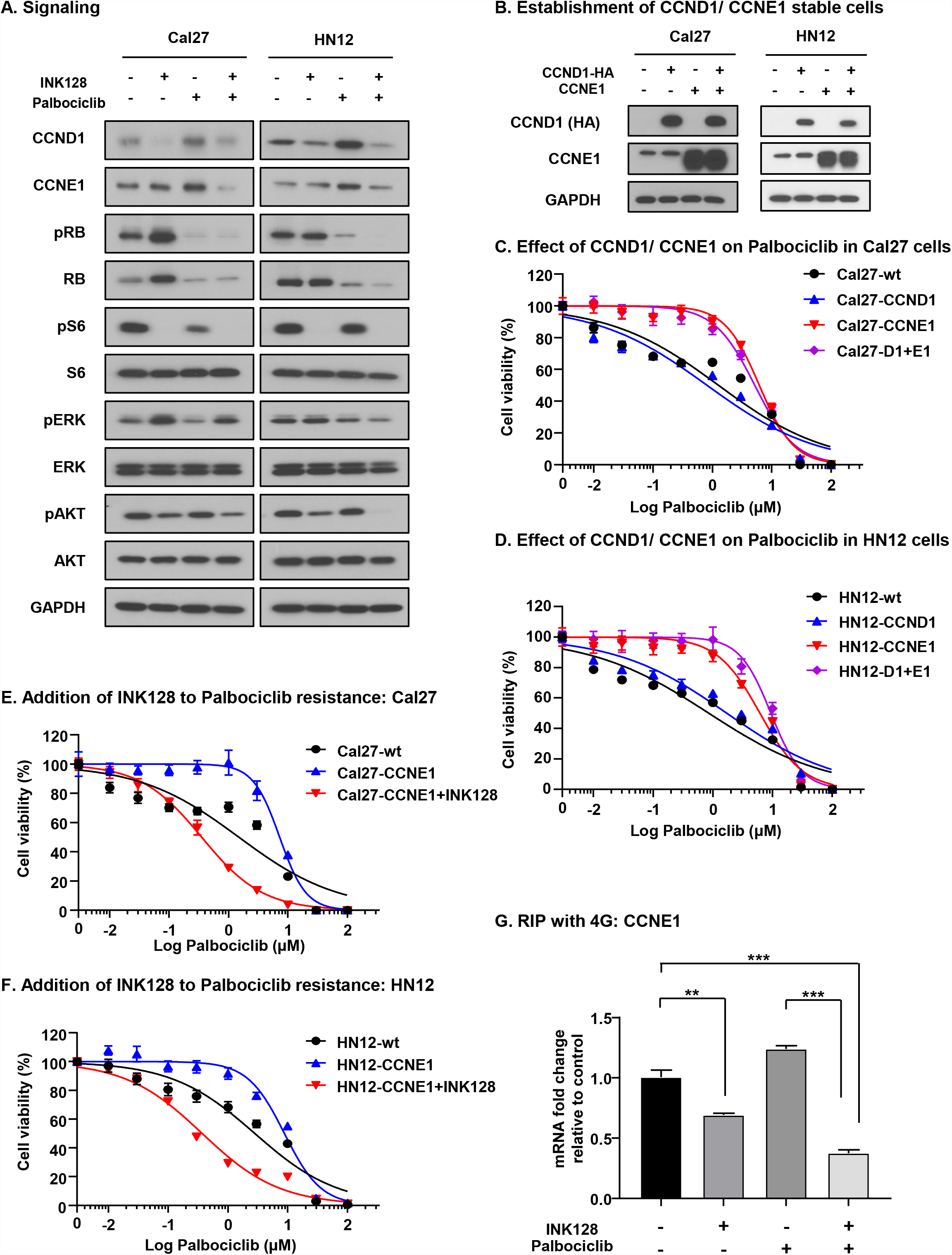
Upregulation of CCNE1 by palbociclib confers resistance to palbociclib, which can be reverted by INK128. A, Signaling change with INK128, palbociclib, and combination. The Cal27/ HN12 HNSCC cells were treated with 50 nM of INK128, 1μM of palbociclib, or both for 2 days after serum starvation overnight, and proteins were subjected to western blot. INK128 could effectively inhibit PI3K/ mTOR activation as judged by pAKT/ pS6 expression. Palbociclib treatment induced upregulation of CCND1 and CCNE1. B, Establishment of stable cell lines with CCND1 and CCNE1 overexpression. Retrovirus with CCND1 was infected to Cal27/ HN12 cells, and lentivirus with CCNE1 was infected to Cal27/ HN12 cells. Cal27-CCND1, Cal27-CCNE1, Cal27-CCND1+CCNE1, HN12-CCND1, HN12-CCNE1, and HN12-CCND1+CCNE1 were established. C, Treatment with palbociclib for Cal27 which stably overexpress CCND1, CCNE1 and both. CCNE1 overexpression was sufficient to induce resistance to palbociclib. Bars represent average ± SEM (N = 3). D, Treatment with palbociclib for HN12 which stably overexpress CCND1, CCNE1 and both. CCNE1 overexpression was sufficient to induce resistance to palbociclib. Bars represent average ± SEM (N = 3). E, Addition of INK128 to palbociclib resistant Cal27-CCNE1 cells. INK128 was added to palbociclib with 1/20 concentration of palbociclib. INK128 completely abolished the resistance to palbociclib in Cal27-CCNE1. Bars represent average ± SEM (N = 3). F, Addition of INK128 to palbociclib resistant HN12-CCNE1 cells. INK128 was added to palbociclib with 1/20 concentration of palbociclib. INK128 completely abolished the resistance to palbociclib in HN12-CCNE1. Bars represent average ± SEM (N = 3). G, eIF-4G binding assay with INK128 or palbociclib treatment for *CCNE1*. INK128 reduced eIF4G binding mRNA of *CCNE1* in Cal27 cells, and combination treatment with INK128 and palbociclib significantly reduced mRNA expression of *CCNE1* binding with eIF4G compared with control or palbociclib treatment. Bars represent average + SEM (N = 3). \*\**P* < 0.01, \*\*\**P* < 0.001 (one-way ANOVA).

### Combination therapy with INK128 and palbociclib is effective against HNSCC xenograft

Next, we asked if this combination of INK128 and palbociclib is effective *in vivo*. Using Cal27 and HN12 xenograft models, we started treatment with INK128, palbociclib, or combination after tumors were established. Since high frequency of myelosuppression has been reported for palbociclib in clinical trials (20), we used relatively low dose palbociclib for this in vivo study (50mg/kg/day). In our Cal27 xenograft model, INK128 or palbociclib treatment as a single agent did not inhibit tumor growth, but the combination of these drugs significantly inhibited tumor growth (Fig. 4A). As for HN12 xenograft, palbociclib did not inhibit tumor growth, and INK128 was relatively effective as a single agent, but combination therapy had significantly stronger effect than single agents (Fig. 4B). The H&E staining of these tumors showed that mTOR inhibition together with palbociclib caused tumor collapse with smallest residual tumor masses at the end of the treatment (Fig. 4C and 4D). To assess the inhibition of proliferation *in vivo*, we used BrdU staining for tumors with short-term treatment of palbociclib, INK128, or combination. The combination therapy demonstrated the lowest percentage of BrdU positive cells in both Cal27 and HN12 xenograft which indicate strong inhibition of cell proliferation in co-administered tumors (Fig. 4E and 4F).

**Figure 4.**
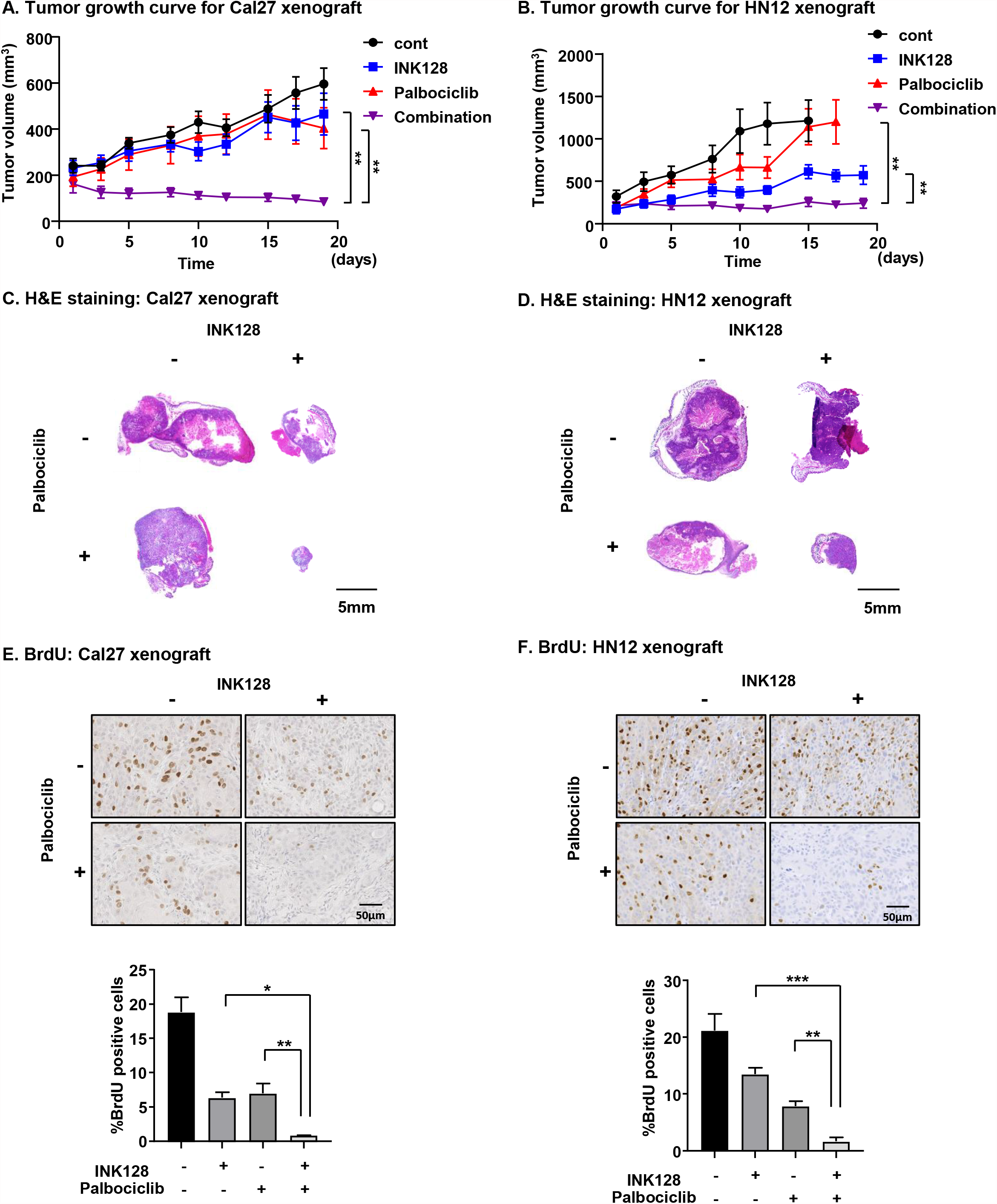
Combination therapy with INK128 and palbociclib is effective against HNSCC xenograft. A, Tumor growth curve for Cal27 xenograft with INK128, palbociclib, and combination. INK128 or palbociclib as a single agent did not inhibit tumor growth of Cal27 xenograft, but combination of these drugs significantly inhibited tumor growth. Bars represent average ± SEM (N = 10). \**P* < 0.05, \*\**P* < 0.01 (unpaired T test at day 18). B, Tumor growth curve for HN12 xenograft with INK128, palbociclib, and combination. Palbociclib did not inhibit tumor growth, and INK128 was relatively effective as a single agent, but combination therapy had significantly stronger effect than single agent. Bars represent average ± SEM (N = 10). \*\**P* < 0.01 (unpaired T test at day 15). C, H&E staining of Cal27 xenograft tumors. D, H&E staining of HN12 xenograft tumors. mTOR inhibition together with palbociclib caused tumor collapse with nearly nonexistent residual tumor masses at the end of the treatment. E, BrdU staining for Cal27 xenograft tumors. After 5 days treatment with INK128, palbociclib, or combination, the tissue was subjected to BrdU staining. The tumor with combination therapy demonstrated significantly fewer percentage of BrdU positive cells compared with the tumors treated with single agent. Bars represent average + SEM (N = 3). F, BrdU staining for HN12 xenograft tumors. The tumor with combination therapy demonstrated significantly fewer percentage of BrdU positive cells compared with the tumors treated with single agent. Bars represent average + SEM (N = 3). \**P* < 0.05, \*\**P* < 0.01, \*\*\**P* < 0.001 (unpaired T test).

## Discussion

The frequent genomic alterations in *CDKN2A* and *CCND1* in HPV-negative clinical HNSCC cases suggest that there is a strong rationale to target CDK4/6 to inhibit tumor progression in HNSCC. Several selective CDK4/6 inhibitors are available in the clinic, such as abemaciclib, ribociclib, and palbociclib. Among them, palbociclib is the first FDA-approved CDK4/6 specific inhibitor, inducing G1 arrest, with a concomitant reduction of phosphorylation of the Rb protein (21). It is approved for advanced or metastatic hormone receptor-positive (HR+) and human epidermal growth factor receptor 2-negative (HER2-) breast cancer (BCa), in combination with endocrine therapy (22,23). For HPV-negative HNSCC, several clinical trials have been conducted using CDK4/6 inhibitors. In selected patients with *CDKN2A*-altered HNSCC, palbociclib monotherapy showed modest antitumor activity (4). Also, in the PALATINUS study, the combination of palbociclib and cetuximab did not prolong the OS in unselected patients (5). To strengthen the anti-tumor activity of palbociclib in HNSCC, novel strategies are needed. In the subgroup analysis in PALATINUS patients, trends for better OS were observed in patients with *CDKN2A* mutations or CDK4/6 amplification, but in the absence of *PIK3CA* alterations (5). Consistent with these data, basic studies showed that *PIK3CA*-mutant HNSCC cells are less responsive to palbociclib (24). These results are consistent with the results of our study showing that mTORi and palbociclib could have beneficial combinatory effects on HNSCC.

The therapeutic potential of mTORi for HNSCC has been extensively studied. Our group pioneered the use of rapamycin as a single agent to treat HNSCC xenograft (6). In this early study, we showed that phosphorylated S6, the most downstream target of the Akt-mTOR pathway, is frequently accumulated in HNSCC clinical specimens. Furthermore, we used rapamycin to treat four different types of HNSCC xenografts, resulting in tumor regression. Following this study, several groups have reported the effectiveness of mTORi for HNSCC (25-27). In turn, these analyses from basic research led to multiple clinical trials including single agent mTORi, or combined treatment with mTORi and other agents (9,28-31). Our group has recently shown the efficacy of rapamycin as monotherapy for previously untreated patients. 21 days treatment for 16 patients with rapamycin resulted in 1 complete response, 3 partial response, and 12 stable disease, supporting the potential role of mTORi for HNSCC (7). Furthermore, clinical trials with administration of metformin, which has been shown to regulate mTOR via AMPK, to premalignant lesions of HNSCC has been conducted, and it shows promising results as judged by pathological responses (32). Similarly, we have recently shown that mTOR inhibition with everolimus in the adjuvant setting after definitive treatment of locally advanced HNSCC lesions reduces significantly tumor relapse, specifically in HPV negative cases (8). However, a clinical trial targeting mTOR in heavily pretreated HNSCC patients did not show clinical benefit with everolimus (9). These findings suggest that previous treatments may cause genetic alterations and epigenetic changes in cancer cells; consequently, more complicated mechanisms driving cell growth may be active in these lesions when compared to the use of mTORi in newly diagnosed HNSCC cases, or as an adjuvant post-surgery and/or radiation. In addition, in these early clinical trials mainly three mTORi were used; rapamycin (sirolimus), everolimus and temsirolimus. These three mTORi are often referred as first generation mTORi, blocking only mTORC1. In our study, we used INK128, which is a second generation mTORi that binds to the ATP-binding site of mTOR and inhibits the catalytic activity of both mTORC1 and mTORC2 without inhibiting other kinases (12). In this regard, INK128 is different from previous mTORi, and the anti-tumor effect of second generation mTORi is promising (33).

To overcome potential mechanisms limiting the response to mTORi, we hypothesized that the administration of mTORi to HNSCC combined with targeting agents suppressing resistance pathways may provide better outcomes. In this study, we applied an unbiased approach to find synthetic lethal and resistance targets for INK128 and showed that the cell cycle pathway can be a synthetic lethal target with INK128. Xenograft experiments using human HNSCC cells showed promising results with the co-administration of INK128 and palbociclib. Mechanistically, we showed that INK128 could inhibit the adaptive accumulation of CCNE1 caused by palbociclib. Since INK128 blocks mTORC1 and mTORC2, it can inhibit phosphorylation of 4E-BP1 strongly, which in turn reactivates the tumor suppressive activity of 4E-BP1 (11,12). Dephosphorylated 4E-BP1 associates with eIF4E, and inhibits binding between eIF4E and eIF4G, resulting in reduced translation of mRNAs that are essential to cell proliferation for tumor (11). In this case, one of the eIF4G-biding mRNAs by reduced INK128 is *CCNE1*. This may explain the reduced level of CCNE1 protein after INK128, and the efficacy of combination therapy with INK128 and palbociclib (see Fig. 5). Aligned with this possibility, recent studies suggest that enhanced *CCNE1* mRNA expression levels in BCa patients are associated with resistance to palbociclib (19,24,34). In our case, CCNE1 overexpression was sufficient to induce palbociclib resistance in HNSCC cells.

**Figure 5.**
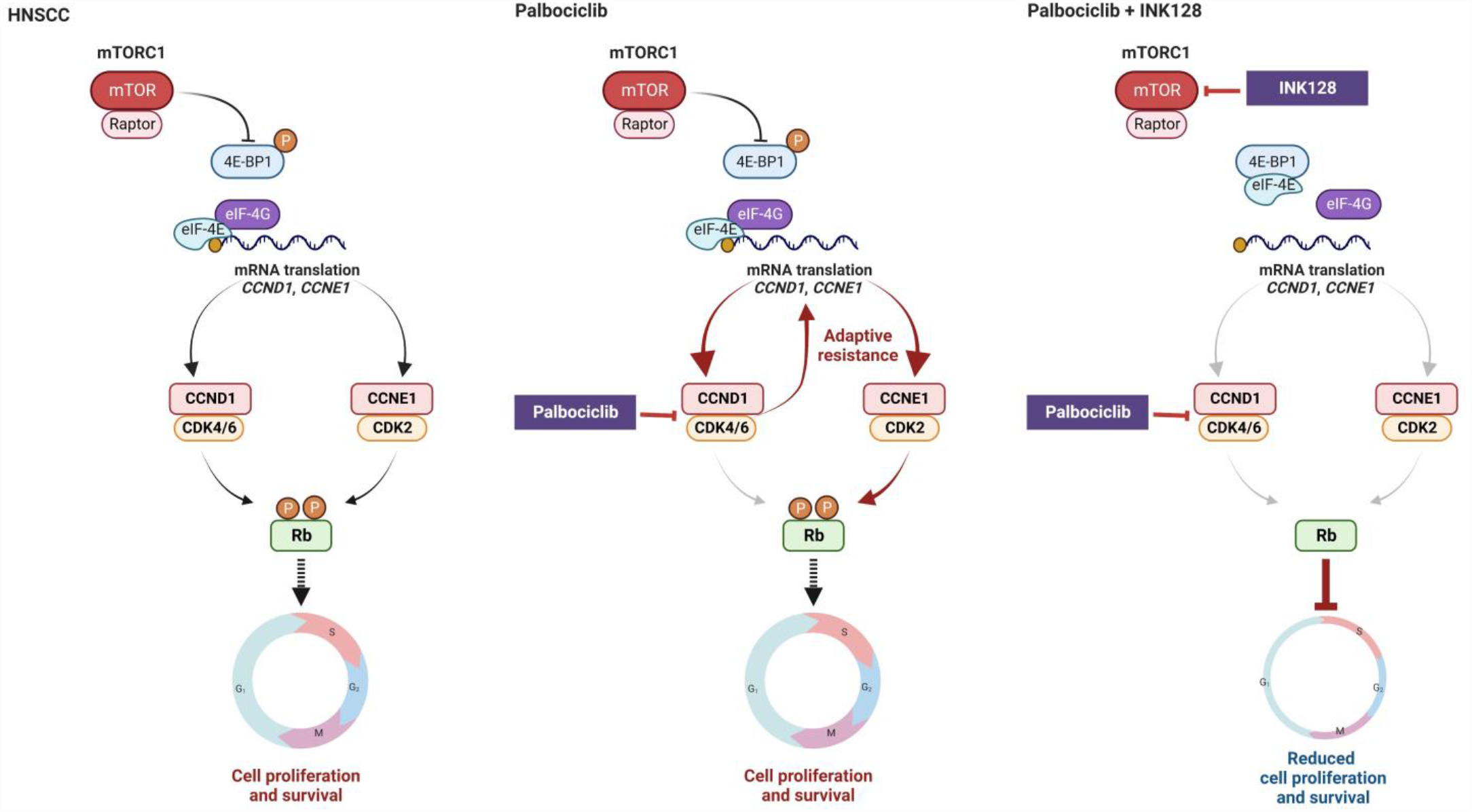
Schematic representation of the mechanism for combination treatment with INK128 and palbociclib. See Discussion for details.

In summary, our unbiased genetic library screen approach revealed that concomitant mTOR blockade reverts the adaptive resistance to palbociclib. Specifically, CCNE1 overexpression caused by palbociclib can be abolished by co-administration of INK128. Ultimately, our findings may provide a novel strategy for HPV negative HNSCC patients by co-targeting mTOR and key cell cycle regulating molecules, which can also have an impact in multiple cancer types that fail to respond to CDK4/6 inhibitors as single agents.

## Materials and methods

Additional information can be found in Supplementary Materials and Methods.

### Cell lines, culture conditions and chemicals

Human HNSCC cell lines HN12 and CAL27 were genetically characterized as part of NIH/NIDCR Oral and Pharyngeal Cancer Branch cell collection, and have been described previously (15).

### CRISPR screen

CRISPR screen was performed as previously described (35,36). Briefly, Cal27-Cas9 cells were infected with Human Kinome CRISPR pooled library (Brunello), and were treated with 2 different groups; vehicle treated or INK128 10nM treated group. The change in the relative abundance of each sgRNA in the library over time is measured using PinAPL-Py software (37). Significantly changed hit sgRNAs were extracted with adjusted p value < 0.001. The hit sgRNAs were subjected to pathway analysis using Enrichr software (38). KEGG pathway combined score was calculated with p-value and z score as follows; c = log (p) * z, where c = the combined score, p = Fisher exact test p-value, and z = z-score (39).

### Antibodies

Antibodies against CCND1 (#2978), CCNE1 (#20808), pRb (#9307), Rb (#9309), pS6 (#2211), S6 (#2217), pERK (#4370), ERK (#9102), pAKT (#4060), AKT (#9272), Cas9 (#14697), HA-Tag (#3724), β-Actin (#4967) and GAPDH (#2118) were purchased from Cell Signaling Technology (Beverly, MA, USA). Antibody against eIF4G (#sc-133155) was purchased from Santa Cruz Biotechnology (Dallas, TX, USA).

### Cell viability assay

Cell viability assay was performed as previously described (40).

### Orosphere assay

Orosphere assay was performed as previously described (32).

### RNA isolation from RNA-binding proteins, polysome analysis, and quantitative PCR

RIP assay was performed following the manufacturer’s instructions (EZ-Magna RIP RNA-binding Protein Immunoprecipitation Kit, Sigma-Aldrich, #17-701). Antibody against eIF4G was used for the part of immunoprecipitation. cDNA synthesis of input RNA and eIF4G binding RNA were followed by qPCR as described before (11).

### Animal work

All the mice studies were approved by the Institutional Animal Care and Use Committee (IACUC), University of California, San Diego (protocol #S15195). The mice were randomized into groups and treated by intraperitoneal injection (ip) with INK128 (1 mg/kg/day, five times a week) or oral gavage with palbociclib (50mg/kg/day, five times a week), or control diluent (10 tumors per each group).

### Tissue analysis

All samples were fixed in zinc formalin (Z-Fix, Anatech) and embedded in paraffin; 5 μm sections were stained with Hematoxylin-Eosin for diagnostic purposes. The immunohistochemistry (IHC) was performed as previously described (11)

### Genomic data analysis

mRNA and RPPA expression analyses where performed using publicly available data generated by The Cancer Gene Atlas consortium, accessed through cBioportal (www.cbioportal.org) (41,42).

## Statistical analysis

All data analysis was performed using GraphPad Prism version 8.02 for Windows (GraphPad Software, San Diego, CA, USA). Comparisons between experimental groups were made using one-way ANOVA or unpaired T test. The overall survival time were assessed using the Kaplan-Meier method and compared using the log-rank test. Asterisks denote statistical significance (non-significant or N.S., *P* > 0.05; \**P* < 0.05; \*\**P* < 0.01; and \*\*\**P* < 0.001). All data are reported as mean ± standard error of the mean (SEM).

## Acknowledgments

Y.G. is supported by the JSPS Overseas Research Fellowships, the Uehara Memorial Foundation Research Fellowship, and the SGH Foundation Cancer Research Fellowship.

Y.G. initiated the study; Y.G. and J.S.G. designed the study and experiments; Y.G., X.W., N.A, K.F., M.A., and A.K. performed the CRISPR screening; Y.G., K.K., T.A., H.I. and X.F. performed *in vitro* experiments, Y.G., K.K., T.A., N.A. and J.S.G. prepared the manuscript, Z.W., N.A., P.M. and J.S.G. provided advice and supervised the project. All authors discussed the results and reviewed the manuscript.

All cartoon renderings were created with the BioRender online platform (BioRender.com).

## Competing Interests

J.S.G. is consultant for Domain Therapeutics, Pangea Therapeutics, and io9, and founder of Kadima Pharmaceuticals, outside the submitted work.

## Figure Legends

**Supplementary Figure 1**. A, Cas9 expression of Cal27 cells. Cas9 expression was confirmed by Western blots. B, Quantification of the AAVS1 locus editing frequency. NGS showed 83.0% and 98.3% of non-homologous end joining (NHEJ) frequency for gT1 and gT2, respectively. C, eIF-4G binding assay with INK128 or palbociclib treatment for *CCND1*. INK128 reduced eIF4G binding mRNA of *CCND1* in Cal27 cells, but combination treatment with INK128 and palbociclib did not change mRNA expression of *CCND1* binding with eIF4G. Bars represent average + SEM (N = 3). \**P* < 0.05 (one-way ANOVA).

**Supplementary Table 1**. Depleted gRNAs in mTORi-treated cells compared with vehicle-treated cells.

