## Supplemental Figure for "A kinome-wide synthetic lethal CRISPR/Cas9 screen reveals that mTOR inhibition prevents adaptive resistance to CDK4/CDK6 blockade in HNSCC"

### Supplementary Figure 1

#### A. Cas9 expression of Cal27 cells

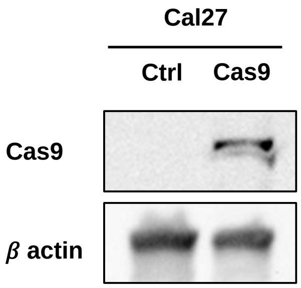


#### B. Quantification of the AAVS1 locus editing frequency

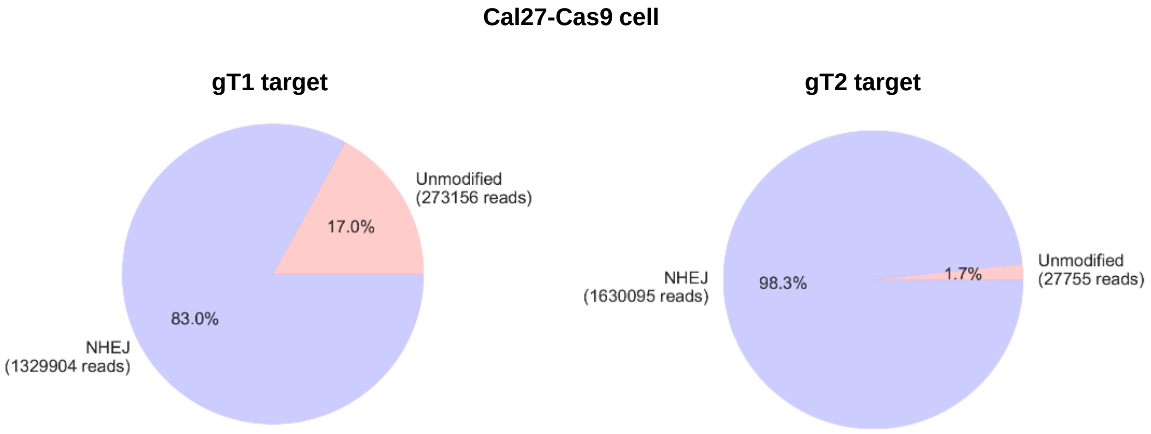


#### C. RIP with 4G: CCND1

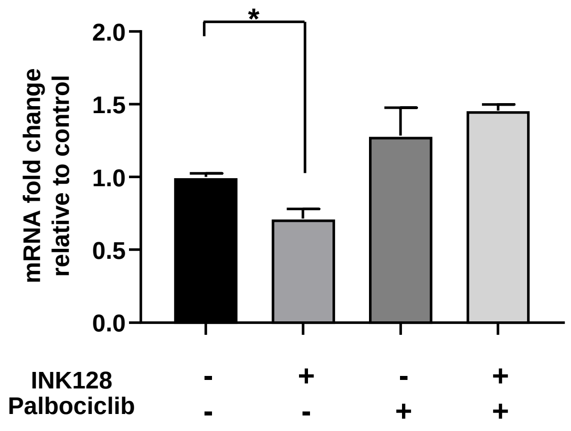
