## Supplemental Materials and methods for "A kinome-wide synthetic lethal CRISPR/Cas9 screen reveals that mTOR inhibition prevents adaptive resistance to CDK4/CDK6 blockade in HNSCC"

### Supplementary Materials and methods

**Cell lines, culture conditions and chemicals.** Human HNSCC cell lines were cultured in DMEM (D-6429, Sigma-Aldrich, St. Louis, MO), 10% fetal bovine serum (FBS) (F2442, Sigma-Aldrich), 1% antibiotic/ antimycotic solution (A5955, Sigma-Aldrich), 5% CO2, at 37°C. INK128 (I-3344) was purchased from LC Laboratories (Woburn, MA), and palbociclib (S1116) was purchased from Selleckchem (Houston, TX).

**CRISPR screen.** The two-vector system was used for this study. First, we generated Cas9-stably expressing Cal27 cells with lentiviral infection from lentiCas9-Blast. lentiCas9-Blast was a gift from Feng Zhang (Addgene plasmid # 52962; http://n2t.net/addgene:52962; RRID: Addgene_52962). The infected cells were selected with blasticidin (10 µg/mL) for 10 days. After confirming Cas9 expression by western blot, the Cal27-Cas9 cells were infected with two different gRNAs targeting AAVS locus, and were subjected to NGS to confirm cutting efficiency. The CRISPR cutting efficiency of Cas9 expressed cells was tested by two AAVS locus targeting sgRNA: gT1, and gT2 (Hygromycin B resistant). AAVS1 gene from genome DNA was amplified and then NEBnext primers (E7335S) were used to attach the sequencing adaptors. Sequencing data were analyzed using CRISPResso2 (http://crispresso.rocks/) [1]. AAVS1 locus targeting sgRNA constructs (gT1/gT2) provided by Prashant's lab. Primer sequences for NGS (Next Generation Sequencing):

NGS_AAVS1_F:
ACACTCTTTCCCTACACGACGCTCTTCCGATCTTCCCAGGGCCGGTTAATGTGG;
NGS_AAVS1_R:
GACTGGAGTTCAGACGTGTGCTCTTCCGATCTTGCCTAACAGGAGGTGGGGGTTAG;

Amplicon reference sequence (+- 30bp around the PAM seq): TGCCTAACAGGAGGTGGGGGTTAGACCCAATATCAGGAGACTAGGAAGGAGGAGGCCTAA GGATGGGGCTTTTCTGTCACCAATCCTGTCCCTAGTGGCCCCACTGTGGGGTGGAGGGG.

Next, Cal27-Cas9 cells were infected with Human Kinome CRISPR pooled library (Brunello) at representation of 650 and a multiplicity of infection (MOI) of 0.3. The viral titer of lentivirus was analyzed using qRT-PCR-titer kit (#631235, Takara, Mountain View, CA), and functional titration. Human Kinome CRISPR pooled library (Brunello) was a gift from John Doench & David Root (Addgene #75312). Cal27-Cas9-kinome library cells were treated with 2 different groups; vehicle treated or INK128 10nM treated group, with triplicate. For PD 0 and PD 20 samples, the barcode was PCR-recovered from genomic samples, and samples where sequenced to calculate abundance of the different sgRNA probes. PCR of sgRNA for Illumina sequencing protocol was obtained from the Broad Institute (https://portals.broadinstitute.org/gpp/public/resources/protocols). The change in the relative abundance of each sgRNA in the library over time is measured using PinAPL-Py software [2]. Significantly changed hit sgRNAs were extracted with adjusted p value < 0.001. The hit sgRNAs were subjected to pathway analysis using Enrichr software [3]. KEGG pathway combined score was calculated with p-value and z score as follows; $c = \log(p) * z$, where c = the combined score, p = Fisher exact test p-value, and z = z-score [4]. Next generation sequencing was conducted by IGM (Institute for Genomic Medicine) Genomics center in UC San Diego.

**Cell viability assay.** 2000 cells were seeded in 96 well plates, and treated as indicated after they attach to the plates. After treatment for 72 hours, culture medium was supplemented with 1/100 of the culture volume of Aquabluer reagent (#6015, MultiTarget Pharmaceuticals LLC, Colorado Springs, CO, USA) for 1h to 4h. Absorbances were recorded at 570 nm in a Biotek Synergy Neo microplate reader. Cell viability assay was performed as previously described [5].

**DNA constructs and viral infection.** pBABE puro cyclinD1 HA was a gift from William Hahn (Addgene plasmid # 9050; http://n2t.net/addgene:9050 ; RRID:Addgene_9050). pInducer20 Cyclin E1 was a gift from Jean Cook (Addgene plasmid # 109348; http://n2t.net/addgene:109348 ; RRID:Addgene_109348). Plasmids were packaged into retrovirus and lentivirus in HEK293T cells respectively, and cells were infected with viruses for 2 days. The infected cells were selected with puromycin (1 μg/mL) for 3 days, or selected with G418 (1000 μg/mL) for 7 days, respectively. To overproduce CCNE1, cells were treated with 1 μg/mL doxycycline for at least 48 hours.

**Orosphere assay.** Cells were seeded in 96-well ultra-low attachment culture plates (Corning, Corning, NY) at 100 cells per well. Medium consisted of DMEM/F12 Glutamax supplement medium (#10565042, Thermo Fisher Scientific), basic fibroblast growth factor (bFGF: 20 ng/ml, #13256029, Thermo Fisher Scientific), epithelial growth factor (EGF: 20 ng/ml, #PHG0313, Thermo Fisher Scientific), B-27 (1:50 dilution, #17504044, Thermo Fisher Scientific), and N2 supplement (1:100 dilution, #17502-048, Thermo Fisher Scientific). Vehicle, INK128 (50 nM), or palbociclib (1 μM) were added when cells were seeded. Around ten days after seeding, photographies were obtained, and the sizes and numbers of sphere colonies on each well were counted using a microscope. This experiment was performed as previously described [6].

**RNA isolation from RNA-binding proteins, polysome analysis, and quantitative PCR.** RIP assay was performed following the manufacturer's instructions (EZ-Magna RIP RNA-binding Protein Immunoprecipitation Kit, Sigma-Aldrich, #17-701). Total RNA was converted to cDNA using SuperScript™ VILO™ cDNA Synthesis Kit (#11754250, ThermoFisher Scientific). qPCR was performed using SYBR™ Select Master Mix (#4472908, ThermoFisher Scientific). The following primers were used for qPCR. CCND1 fwd 5'- AGCTGTGCATCTACACCGAC, CCND1 rev5'- GAAATCGTGCGGGGTCATTG, CCNE1 fwd5'- CCATCATGCCGAGGGAGC, CCNE1 rev5'- GGTCACGTTTGCCTTCCTCT.

**Western blotting.** Exponentially growing cells were washed in cold PBS, lysed on ice in lysis buffer (50 mM Tris-HCl, 150 mM NaCl, 1 mM EDTA, 1% NP-40, supplemented with Halt™ Protease and Phosphatase Inhibitor Cocktail (#78440, ThermoFisher Scientific). Cell extracts were collected, sonicated, and centrifuged to remove the cellular debris. Supernatants containing the solubilized proteins were quantified using the detergent compatible DC protein assay kit (#5000111, Bio-Rad, Hercules, CA, USA). Equal amounts of protein were separated by SDS-PAGE, and transferred to PVDF membranes. For immunodetection, membranes were blocked for 20 min at room temperature in 5% non-fat dry milk in TBST buffer, followed by 2h incubation with the appropriate antibodies, in 3% BSA-T-TBS buffer. Detection was conducted by incubating the membranes with horseradish peroxidase–conjugated goat anti-rabbit IgG secondary antibody (Southern Biotech, Birmingham, AL, USA) at a dilution of 1:20,000 in 5% milk-T-TBS buffer, at room temperature for 40 min, and visualized with Immobilon Western Chemiluminescent HRP Substrate (EMD Millipore, Burlington, MA, USA).

**Animal work.** All the mice studies were approved by the Institutional Animal Care and Use Committee (IACUC), University of California, San Diego (protocol #S15195). To establish tumor xenografts, $2.0 \times 10^6$ cells were transplanted into the flanks of athymic nude mice (female, four to six weeks old) (Charles River Laboratories, Wilmington, MA), and when the tumor volume reached approximately 200 $mm^3$, the mice were randomized into groups and treated by intraperitoneal injection (ip) with INK128 (also known as MLN-0128 or TAK-228, 1 mg/kg/day, five times a week) or oral gavage with palbociclib (50mg/kg/day, five times a week), or control diluent (10 tumors per each group). Tumor volume was calculated by using the formula length × width × width/2. The mice were euthanized at the indicated time points and tumors isolated for histologic and immunohistochemical evaluation.

**Tissue analysis.** All samples were fixed in zinc formalin (Z-Fix, Anatech) and embedded in paraffin; 5 μm sections were stained with Hematoxylin-Eosin for diagnostic purposes. For immunohistochemistry (IHC) studies, the sections were deparaffinized, and hydrated through graded ethanols. The slides were extensively washed with distilled water and antigen retrieval was performed by high temperature treatment with 10 mM citric acid in a microwave. After washing with water and PBS, the slides were successively incubated with the primary and secondary antibodies, and the ABC reagent (Vector Laboratories, Burlingame, CA). The reaction was developed with 3-3'-diamonobenzifdine under microscopic control.
